# Lessons learned from manual curation of thousands of gene models in the nematode *Pristionchus pacificus*

**DOI:** 10.64898/2026.02.18.706511

**Authors:** Christian Rödelsperger, Neha Agyal, Shiela Pearl Quiobe, Hao Wu, Waltraud Röseler, Dafne Ibarra-Morales, Ralf J. Sommer

## Abstract

Continuous developments in sequencing technologies have led to the generation of chromosome-scale genome assemblies across the whole tree of life, but our ability to annotate genomes has lacked behind. One major problem consists in the fact that typically not all genes are expressed at detectable levels at any given life stage or environment. Therefore, available transcriptome data needs to be complemented by gene prediction programs and protein homology evidence. However, how to optimally combine these different data types is not well understood. Here, we present a case study, where we community curated gene annotations of the *Pristionchus pacificus* strain RSC011. By incorporation of new Iso-seq and RNA-seq data and genome-wide screening, we identified and corrected more than 7,500 (∼24%) gene models. While the improved gene annotation for the RSC011 strain will be useful for the *P. pacificus* community, our study reveals several gene annotation problems that may affect data from other species. Among these, we identified assembly errors, artificial transcript fusions resulting from overlapping genes and polycistronic RNAs, falsely called open reading frames, and error propagation based on homology data as frequent sources of gene annotation errors. Thus, our findings may be helpful in guiding future efforts to annotate genomes across different taxonomic groups.

## Introduction

The nematode *Pristionchus pacificus* is an established model for investigating developmental plasticity [1,2], host microbe interactions [3,4], and behavior [5,6]. It complements *Caenorhabditis elegans* by virtue of additional ecological and behavioral traits—most notably, a feeding-structure polyphenism in which individuals adopt either a narrow, bacteriovorous stenostomatous form or a wide, predatory eurystomatous form [7,8]. This discrete mouth form switch, conditioned by a combination of environmental cues and genetic regulators, makes *P. pacificus* especially amenable for dissecting how organisms sense and integrate environmental signals resulting in developmental decisions [2,9].

*P. pacificus* is a cosmopolitan species and ongoing field trips have collected more than one thousand natural isolates of which hundreds have available whole genome resequencing data [10–12]. Among the diverse natural isolates of *P. pacificus*, the strain RSC011 is of particular interest. In contrast to many strains that are biased toward the eurystomatous morph under laboratory conditions, RSC011 maintains a relatively high baseline frequency of the stenostomatous form [7,13]. Previously, we exploited this feature in order to dissect the genetic basis of natural variation in the mouth form plasticity [13]. In that study, we crossed strains differing in mouth form frequencies, created recombinant inbred lines, and performed QTL mapping. This identified a multigene locus including the key regulator *eud-1* whose cis-regulatory architecture was found to modulate the frequency of the predatory morph [13]. More recently, we leveraged the *P. pacificus* RSC011 strain in an experimental evolution study to explore environmental influences on mouth form plasticity and associated transgenerational epigenetic inheritance [14]. Using long-term environmental induction and food-reversal experiments, we observed the dietary induction and subsequent transgenerational inheritance of the eurystomatous morph. By forward genetic screening, we identified a role of ubiquitin ligase EBAX-1/ZSWIM8 in this process. *Ppa-ebax-1* mutants are transgenerational inheritance defective, and *Ppa-*EBAX-1 regulates a microRNA cluster of the family *miR-2235a/miR-35*. One byproduct of that work was the assembly and annotation of the RSC011 genome, which provided a critical resource for identifying regulators of transgenerational inheritance [14]. The RSC011 genome could be assembled and scaffolded to the level of chromosomes and evidence-based gene models were generated by the PPCAC pipeline (‘PPCAC1’, Table 1) that uses transcriptomic and homology data to guide the annotation process [15]. However, the quality of these automatically generated gene annotations is likely inferior to the gene annotations of the commonly used *P. pacificus* ‘wildtype’ reference strain PS312, which had been subject to two rounds of extensive community curation [16,17]. In these two studies, we manually inspected thousands of suspicious candidate loci and could identify and correct artificial gene fusions and add missing genes [16,17]. Similar approaches were more recently adapted by different groups in order to improve the quality of gene annotations of other nematode species [18–21]. Here, we report a community curation effort to improve the quality of the RSC011 gene annotations. This will make the RSC011 genome a better resource for future experimental studies and at the same time, our findings may be used to guide and improve gene annotation processes for other genomes.

**Table 1.** Comparison of different gene annotations.

| Gene annotation set | Number of gene models | Number of problematic models | BUSCO (C[S,D]/F/M) % | with starting Methionine | with annotated 3'UTR |
| --- | --- | --- | --- | --- | --- |
| PPCAC1 | 30,074 | 2138 | 81.5[79.0,2.5]/1.9/16.6 | 14,016 | 3156 |
| PPCAC1<br>polished | 31,701 | 1114 | 87.4[84.5/2.9]/1.2/11.4 | 14,231 | 2019 |
| PPCAC2<br>vanilla | 31,726 | 2819 | 86.9[84.1/2.9]/1.6/11.5 | 22,921 | 16,049 |
| PPCAC2<br>cream | 29,497 | 697 | 88.6[85.6/3.0]/1.2/10.2 | 22,096 | 16,265 |

The table summarizes features of different gene annotations for the *P. pacificus* strains RSC011. The gene set 'PPCAC1' denotes the previously published gene annotation [14]. The gene set 'PPCAC1 polished' represents the reannotation of the polished genome assembly using the same protocol and data sets as in Quiobe et al. [14]. 'PPCAC2 vanilla' denotes a fully automated gene annotation that incorporates newly generated RNA-seq and Iso-seq data. 'PPCAC2 cream' denotes the final manually curated gene annotation. Problematic gene models were defined as gene models with three or more 3'UTR or 5'UTR exons. Completeness values were calculated using BUSCO (version 5) against the nematode odb10 data set. Note that the number of missing BUSCO genes is an overestimate as it has been demonstrated that extensive sequence divergence can result in a failure to detect orthologs [15].

## Results

### Retained introns, overlapping genes, polycistronic reads, and assembly errors cause gene models with unusually long UTRs

In order to improve the gene annotations for the *P. pacificus* strain RSC011, we generated Iso-seq data of mixed-stage populations and RNA-seq data of different developmental stages (eggs, J2, J3, J4, and adult worms). All transcriptomic data were generated using nematode cultures that were grown on *E. coli* OP50 bacteria as diet. Overall, we obtained 1.37 million HiFi reads with a median length of 2.1 kb. These Iso-seq reads were aligned against the RSC011 genome assembly by the software GMAP and then assembled by the software stringtie [22,23]. Illumina RNA-seq data was first assembled into longer transcripts using the software trinity [24], and then processed in the same way as the Iso-seq data. We adapted the PPCAC annotation pipeline [15] to incorporate these transcriptomic gene models (Fig 1). Thereby, we noticed that 4113 gene models had more than three exons that were annotated as untranslated regions (UTR) either as 3’ or 5’. Following previously established protocols for community curation, we inspected several hundreds of such cases manually in the genome browser [16,17,25]. This showed that those unusually long UTRs were frequently caused by retained introns, overlapping genes, polycistronic reads, and assembly errors (Fig 2A-C). Note that nematodes have operon-like structures that produce polycistronic transcripts which are later processed into mature moncistronic transcripts by trans-plicing [19]. Most cases of polycistronic reads in the RSC011 genome appear to be derived from such immature operon transcripts. While in many cases, community curation was able to replace problematic cases with alternative gene models, this method could not easily correct misannotations resulting from assembly errors. Therefore, we decided to create a polished version of the RSC011 genome.

**Figure 1.**
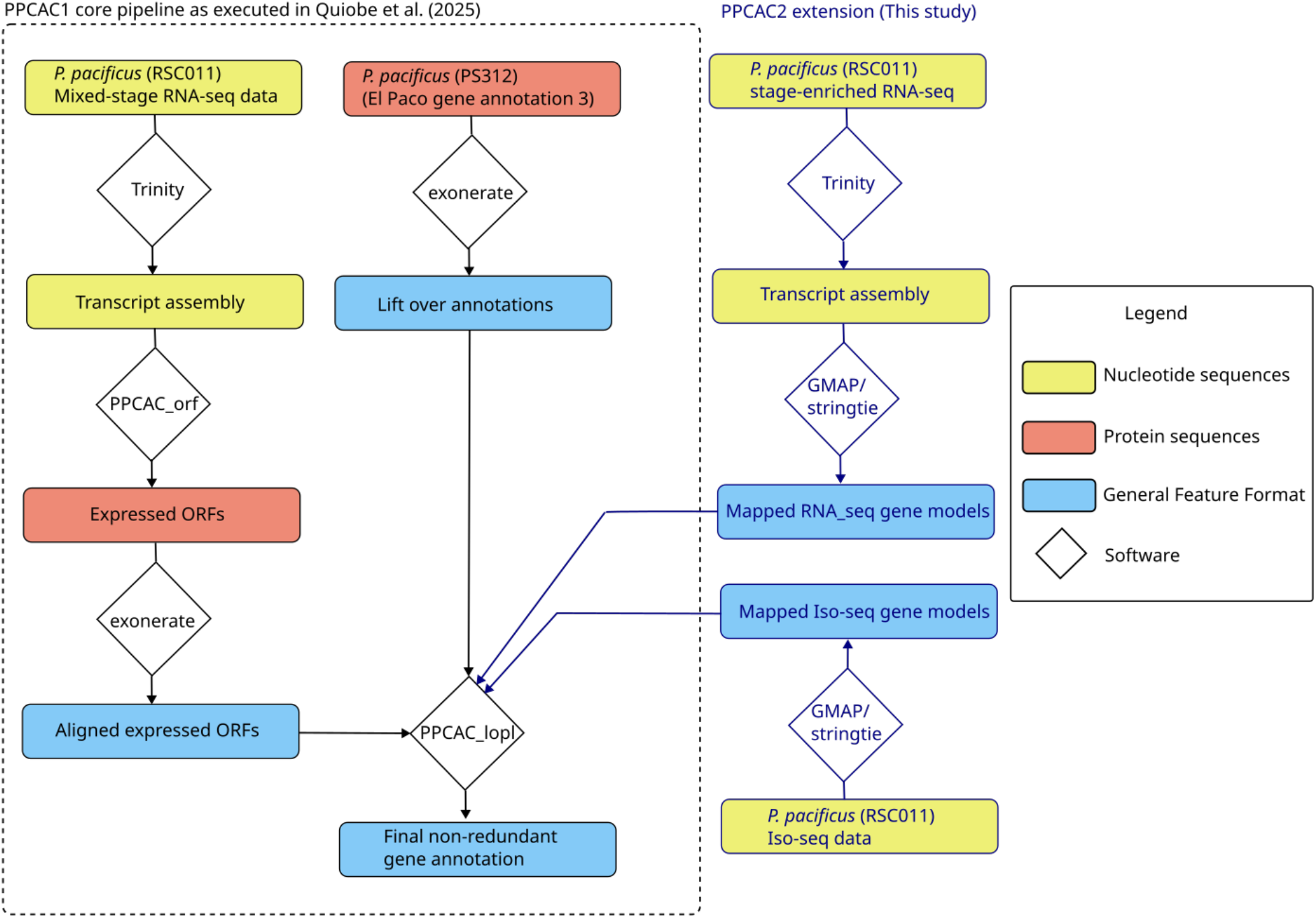
Gene annotation workflow. The flow diagram shows the simplified workflow for annotating the *P. pacificus* RSC011 genome using the PPCAC pipeline [14,15]. For this study, we extended this workflow in order to integrate newly generated Iso-seq and RNA-seq data.

**Figure 2.**
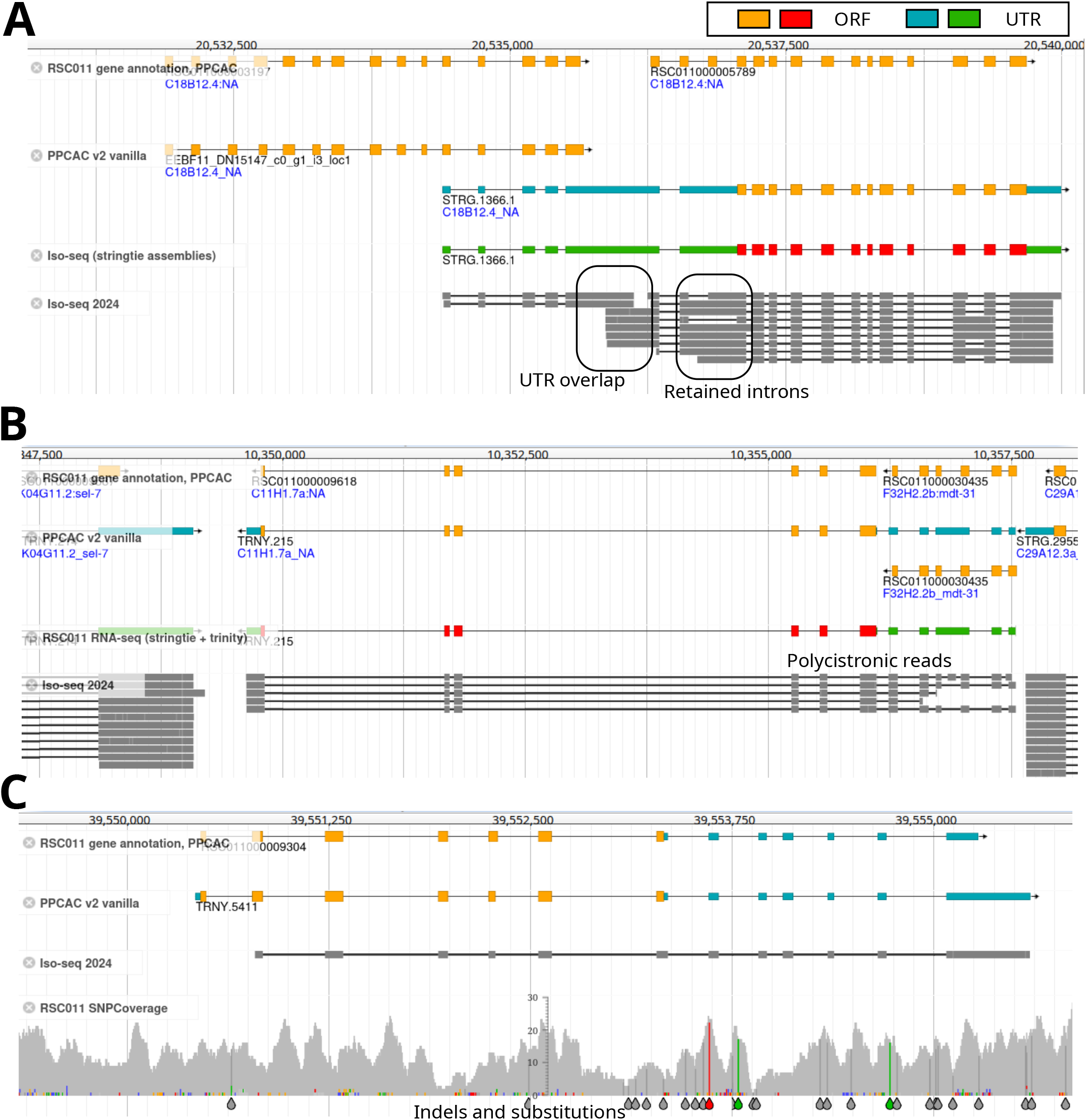
Retained introns, overlapping genes, polycistronic reads, and assembly errors cause gene models with unusually long UTRs. **A)** The genome browser screenshot shows a roughly 10 kb region in the RSC011 genome assembly with the original annotation as topmost track [14]. BLASTP searches of the two paralogous genes against *C. elegans* support that these are two separate genes since both copies are homologous to the same part of the C. elegans gene *C18B12*.*4*. The new annotation (PPCAC v2), the raw Iso-seq alignments together with the assembled transcripts are shown in separate tracks. While both copies are at least partially covered by Iso-seq reads, both genes appear to overlap in the UTR sequences. Together with frequently retained introns, this leads to a transcript assembly with unusually long 5’ UTR sequence (STRG.1366.1). **B)** This genomic region encodes two genes that are homologous to two different *C. elegans* genes. However, some polycistronic Iso-seq reads span both genic loci. **C)** This genome browser screenshot shows a gene model with a long UTR sequence. The additional RSC011 SNPCoverage track summarizes whole genome resequencing data at this locus and reveals SNPs and indels that likely introduce frameshifts and early stop in the ORF.

### Genome assembly polishing greatly reduced the number of problematic gene models

To correct potential assembly errors in the RSC011 genome, we leveraged existing whole genome sequencing data from over 160 mutant lines that were derived from the RSC011 strain [14] and searched for substitutions and indels that were found in more than 100 of these mutant lines. This resulted in 1883 substitutions and 42,138 indels that likely represented assembly errors. We generated a polished version of the RSC011 genome assembly using the software mutsim [26]. Generating gene annotations for the polished genome assembly using the same computational procedures as described above (‘PPCAC2 vanilla’, Table1) reduced the numbers of problematic gene models (gene models with either more than three 3’UTR or 5’UTR exons) from 4113 to 2819. This reduction in the number of problematic gene models was also observed when running the original annotation procedure as described previously [14,15] on the polished genome (‘PPCAC1 polished’, Table 1). In addition, both sets of annotations showed a pronounced increase in the completeness value (Table 1), suggesting that even genome assemblies that were generated using HiFi reads may still contain substantial numbers of assembly errors. These errors will affect over one thousand gene models and additional genome assembly polishing can thus also improve the quality of gene annotations.

### Community curation corrects more than two thousand problematic gene models

In order to correct the remaining problematic gene models, we employed a community annotation effort where the list of 2819 problematic gene models were shared as an online spreadsheet among four community curators who inspected these cases in the genome browser and selected alternative gene models from various computationally derived models (RNA-seq gene models, Iso-seq gene models, RNA-seq derived ORFs only, gene models with truncated UTRs). This approach is different from platforms such as web apollo [27], which allow very flexible modifications of individual gene models. Only allowing curators to choose among predefined gene models, requires less training and makes this approach more scalable to larger numbers of candidate sets. In hindsight, many cases will remain uncorrected as the predefined gene models often do not allow a satisfiable solution. For those cases, the curators could note this as an additional comment. In most cases, the curators had to frequently decide whether a locus encoded two genes or one. Two or more genes can be merged due to overlapping transcribed loci (Fig 2A,B) or due to incorrectly predicted gene boundaries in homology derived models. During the community curation, we used clearly separated transcriptomic alignments between two neighboring gene models or opposite strand orientation with additional conservation in *C. elegans* (BLASTP e-value < 0.001) as strong indicators supporting the presence of two genes. Furthermore overlapping ORFs, RNA-seq coverage differences, and conservation in *C. elegans* were used as weaker evidence to support either the presence of one or two genes at a locus. After manual inspection and correction of all 2819 problematic gene models, the number of genes with more than three UTR exons could be reduced to 697. This confirms our previous experience that community-based curations are an effective means to reduce the number of problematic gene models [16,17].

### Genome-wide screens identify roughly 24% of gene models as problematic

During the community curations, we occasionally observed erroneous annotations of neighboring genes (e.g. artificially fused genes not resulting in multiple UTR exons, predicted antisense ORFs, homology derived gene models with neither transcriptomic support nor conservation in *C. elegans*). To screen and correct additional gene annotation errors, we scanned the complete RSC011 assembly in the genome browser at 20 kb resolution. This also allowed us to recheck and eventually modify curations from the first round. For the whole genome, this resulted in a total of 7528 corrections (S1 Table). Automated classification of these changes yielded 3562 removals, 2928 single gene replacements, 835 splits, and 204 added genes (Fig 3A). Note that some of those cases actually represent gene fusions (Fig 3B), but in the shared spreadsheet, such fusions were recorded as two independent corrections: one replacement and one removal. Overall, these results suggest that roughly 24% of the 31,726 automatically annotated gene models have obvious problems that could be easily detected by visual inspection in the genome browser.

**Figure 3.**
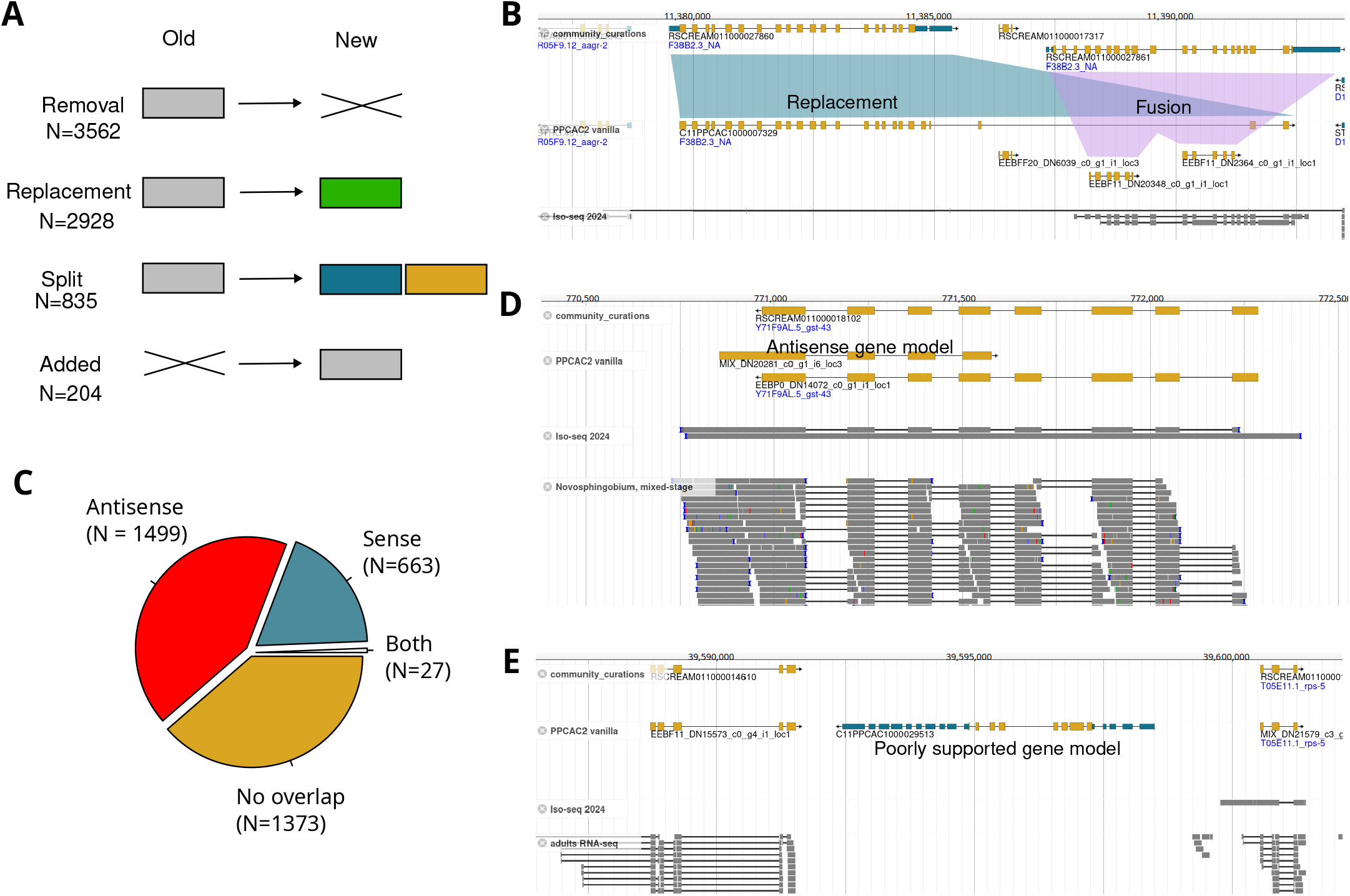
Roughly 24% of gene models are problematic. **A)** Genome-wide screening for obvious gene annotation artifacts led to over 7500 proposed corrections. Those corrections were classified into replacements, splits, removals and additions. **B)** At this locus, C11PPCAC1000027860 was chosen initially as the gene model with the longest ORF. Subsequently, only smaller gene models were added in order to prevent overlaps. However, transcriptomic data supports two genes at this locus. This could be corrected by one replacement and one fusion. **C)** Removed genes were further subgrouped based on overlaps with the final community curated gene annotations. The largest group consisted of antisense gene models and poorly supported gene models without any overlaps. **D)** This genome browser screenshot shows an example of an antisense gene model that has very similar gene structure with a gene on the opposite strand that encodes a protein with a homolog in *C. elegans*. **E)** This region shows a poorly supported gene model that was removed after community curation.

### Antisense coding potential frequently causes misannotated ORFs

The largest class of corrections corresponds to removed gene models. We subdivided this class based on exonic overlaps with gene models in our final gene annotation (‘PPCAC2 cream’, Table 1). Notably, the largest group consists of removed genes that overlap with protein coding exons on the opposite strand (Fig 3C,D). In many of those cases, one of the overlapping gene models shows conservation with *C. elegans*, suggesting that the antisense gene model is a result of misassigning the ORF to the wrong strand. In many cases, sense and antisense models share the same exon-intron structure suggesting that coding sequences frequently have antisense coding potential, which can lead to incorrectly annotated ORFs. The second most abundant group of removed genes consists mostly of gene models that do not have homologs in *C. elegans* and show very little expression evidence in our transcriptomic data sets (Fig 3E). Many of those poorly supported gene models were already captured in the candidate set with multiple UTR exons as they are unlikely functional and intraspecies variation frequently introduces premature stops during the liftover of gene models from the *P. pacificus* reference strain PS312. Note that removing gene models based on the absence of transcriptomic inherently comes with the caveat that we can never be absolutely sure that a locus does not encode a gene. Eventually, such models will be reactivated if future studies provide more robust evidence that they are largely correct.

Among the removed gene models with same strand overlaps, we found many partial ORFs or isoforms that are part of other better supported gene models. However, such cases are far less abundant than the spurious antisense gene models. We encountered the problem of spurious antisense gene models already previously and therefore incorporated the removal of species-specific orphan genes with antisense overlaps in the annotation process for different *P. pacificus* strain genomes [17,28,29], but it appears that antisense coding potential is not only generating species-specific orphan genes, but can also create gene models with apparent protein conservation across species. This problem could in future be alleviated by incorporating homology information in the ORF prediction or by only using strand-specific RNA-seq data.

### Heuristics favoring longer genes cause propagation of gene fusions errors

One of the most abundant gene annotation problems in the genome of the *P. pacificus* reference strain PS312 consisted in artificial gene fusions that likely resulted from failure of gene prediction programs in identifying the exact gene boundaries [16]. While we have corrected thousands of those instances in our latest set of *P. pacificus* PS312 gene annotations (El Paco gene annotation 3) [16,17], it cannot be excluded that this set of gene annotation still harbors some artificially fused gene models [30]. Using this gene set as homology data to annotate the genome assemblies of other *P. pacificus* strains and other *Pristionchus* species may lead to propagation of these gene annotation errors, especially, when combined with a heuristic to choose the longest ORF per locus as representative gene model [15]. This likely explains the presence of 835 corrections that were classified as splits (Fig 3A). However, we need to emphasize here that the use of the El Paco annotations as homology data is still needed to annotate genes without any expression evidence (Fig 4A) or genes where transcriptomic data is not sufficient to generate a complete gene model (Fig 4B). Thus, future annotation pipelines need to be optimized in how they combine transcriptomic and protein homology evidence. Alternatively, an even better reference data set might prevent further error propagation issues.

**Figure 4.**
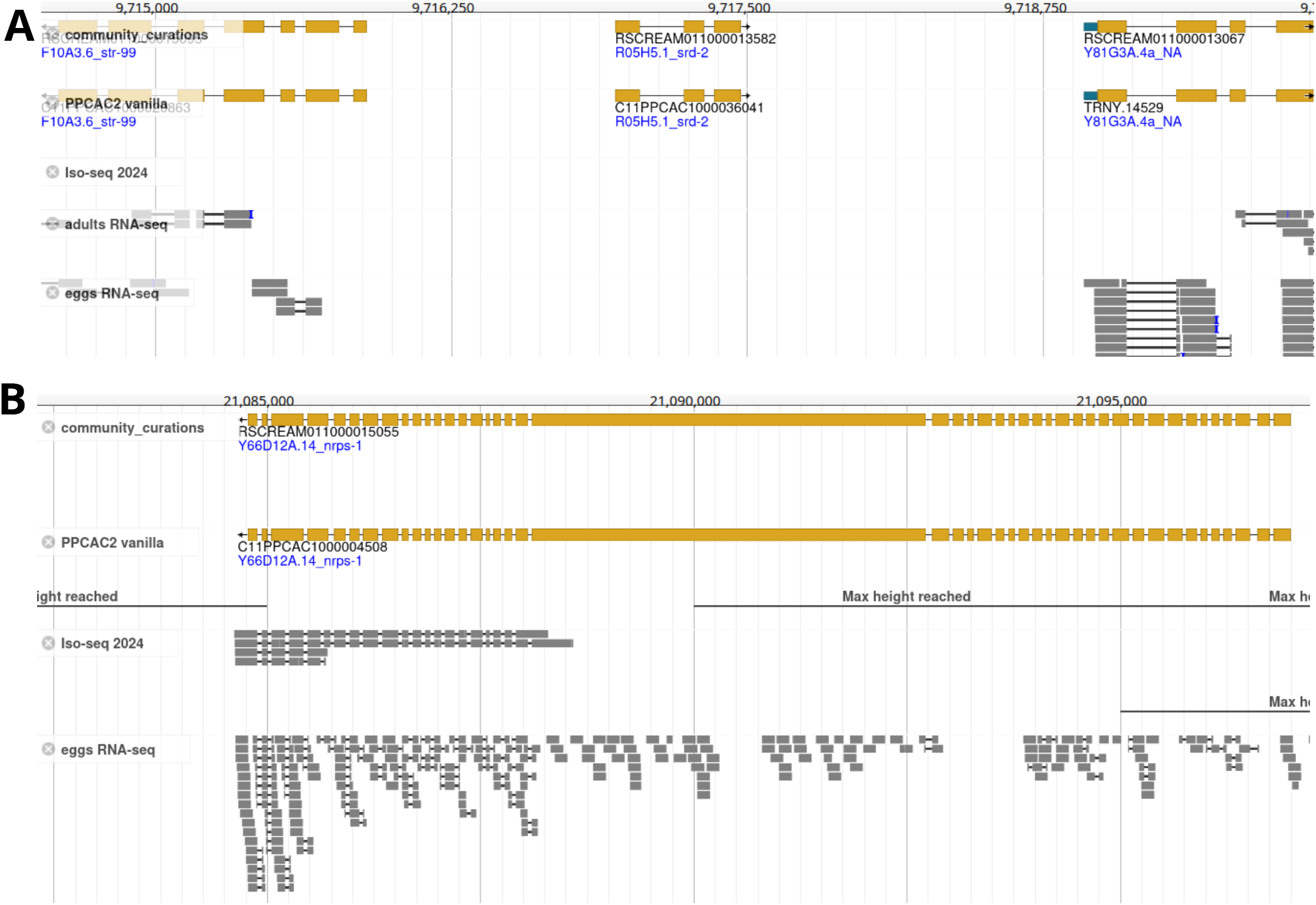
Homology data and gene prediction program help to annotate genes without complete transcriptomic support. **A)** This genomic region shows an example of a gene that has no transcriptomic support in our data, but has a homolog in *C. elegans*. **B)** This genomic locus shows a gene with nearly complete homology in *C. elegans*, but without full transcriptomic support.

### The community curated RSC011 gene annotations exhibit the highest level of completeness

The primary focus of the previous RSC011 gene annotation was to identify protein-coding genes. Given that many assembled transcripts might be incomplete, ORFs were sometimes extended over the starting methionine as long as no inframe stop codon were encountered [14,15]. For this reason, many protein sequences did not start with methionine. In addition, generally no UTR regions were annotated. With the additional transcriptomic data generated for this study, we incorporated many more complete gene models, which should facilitate a better annotation of the starting methionine and the 3’UTR. To systematically compare the final community curated gene annotation with different automatically generated gene annotations, we calculated the number of genes with annotated 3’UTRs, the number of problematic gene models, the number of protein products that start with a methionine, and the completeness value as estimated by the benchmarking of universally single-copy orthologs (BUSCO) approach (Table 1) [31]. This showed that polishing the genome assembly led to a pronounced reduction in the number of problematic gene models and a substantial increase in the BUSCO completeness. In comparison, the effect of community curations on the BUSCO completeness was rather minor. Furthermore, incorporation of additional transcriptomic data and restricting the ORF prediction to complete ORFs drastically increased the number of gene models with a starting methionine and a 3’UTR. At the same time, this additional transcriptomic data increases the chance that an assembled transcript will contain a retained intron. Also, prioritizing ORFs with a starting methionine and an annotated 3’ UTR can sometimes lead to the selection of genes with shorter ORFs.

These two problems largely explain the slight decrease in BUSCO completeness between ‘PPCAC1 polished’ and ‘PPCAC2 vanilla’ (Table 1). Thus, despite a reduction in gene count by more than two thousand, the community curated RSC011 gene annotations shows the highest level of completeness.

## Discussion

In this study, we present an improved version of the genome for the *P. pacificus* strain RSC011. The improvements were achieved through multiple steps, such as genome assembly polishing, incorporation of additional transcriptomic resources, and manual curation. The fact that even after the first two steps of improvement, manual inspection identified around 24% of gene models as problematic, clearly highlights the need for manual curation. The resulting improved gene annotations will be an essential resource for future genomic and experimental research on transgenerational epigenetic inheritance [14]. In addition, this work represents a case study summarizing problems in gene annotation and might be useful for developers and users of gene annotation pipelines [32–38]. As such our study highlights several problems that may impact gene annotation. First, genome assembly polishing is an important first step to improve gene annotation quality. Second, problems in transcriptome assembly may arise due to incompletely spliced reads, polycistronic transcripts, and overlapping genes. Such problems could be partially mitigated by the use of RNA-seq read coverage and homology data to separate such fusions at the transcript level. Third, antisense coding potential may cause ORFs to be misassigned to the wrong strand. This can be alleviated by the use of homology data during ORF prediction or strand-specific RNA-seq data. In addition, the ORF prediction should incorporate the possibility that a transcript could encode multiple ORFs. Fourth, it is not totally clear how to best integrate homology and transcriptomic data. While homology data can cause error propagation problems it may be helpful to fully annotate genes with no or only incomplete transcriptomic evidence. Finally, in the context of antisense transcripts, we saw a lot of transcripts where exons are located in introns of genes on the opposite strand, but encountered very little evidence for antisense overlap between exonic regions. This is consistent with previous analysis of strand-specific RNA-seq data in *P. pacificus* [39] and suggests that at least for *Pristionchus* species it should be legitimate to prohibit antisense exonic overlaps during the gene annotation process.

Notably, many if not all of our findings are not new. The difficulty of gene annotation and the problem of error propagation has been acknowledged previously [40]. Genome assembly polishing has been routinely applied to improve the quality of newly assembled genomes [41,42]. Also, the presence of polycistronic transcripts among Iso-seq reads [43] and spurious antisense gene models was known before [17]. Thus, it could have been possible to anticipate each of these problems in advance without the need to screen a whole genome for annotation artifacts and manually correct thousands of gene models. Confucius once said: “By three methods we may learn wisdom: First, by reflection, which is noblest; Second, by imitation, which is easiest; and third by experience, which is the bitterest.” However, we consider as a major outcome of this study that we bring together all these different findings in a single study and highlight their impact on gene annotation. We hope that this knowledge will be helpful for future efforts in annotating genomes across the tree of life.

## Methods

### Worm culturing and transcriptome sequencing

In order to obtain stage-enriched transcriptomes for the *P. pacificus* strain RSC011, we prepared initial cultures by putting four J4 worms onto each of five 6cm plates with 300µl of *E. coli* OP50. After nine days the plates were full of eggs. We washed one of those plates with M9 to obtain a transcriptome representing mostly eggs. The other plates were grown for an additional day. Here two plates were washed with M9 for the J2 stage. The remaining two plates were washed using M9 into 15ml Falcon tubes and let stand for 5 min. Supernatant containing mostly J2 worms were distributed to fresh 6cm plates with 300µl OP50. After 24h we collected developmental stage J3, after 48h J4 and after 60h young Adult. The pellets of each developmental stage were frozen at -80°C until further use. RNA was extracted using the Zymo Direct-zol RNA Miniprep Kit. RNA was sequenced with Novogene. For the Iso-seq experiment worm pellets were prepared from mixed-stage cultures and RNA was extracted with the Zymo Direct-zol RNA Miniprep Kit. We then quantified the total RNA and quality-controlled it using the Bioanalyzer® RNA Pico Kit. All processed samples had a RIN number >9.0. We prepared Iso-Seq libraries using the following protocol: “Procedure & Checklist - Preparing Iso-Seq® libraries using SMRTbell® prep kit 3.0 (PN 102-396-000 REV02 APR2022)”. We used 500ng of total RNA as input amount and performed duplicates of each sample to obtain enough cDNA. Briefly: we synthesize cDNA according to the protocol above and amplified it with 12 PCR cycles. We pooled together the duplicates of the same sample. We indexed individual libraries using SMRTbell barcoded adaptors during the adapter ligation step of the library prep protocol. After the indexed libraries were quality-controlled using the Bioanalyzer® High Sensitivity DNA kit, we pooled the libraries by combining an equal mass of each barcoded library. We sequenced all the libraries on a Sequel® II system, using the Sequel II Sequencing Plate 2.0 and the Sequel II binding kit 3.1, which is recommended for libraries with fragments shorter than 3 kb.

### Transcriptome analysis

From the raw PacBio reads, consensus HiFi reads were generated with the ccs software (version: 6.4.0 with –min-rq 0.99 option) and demultiplexed by lima (version: 2.7.1 with – split-bam-named –same –dump-removed options). HiFi reads were then aligned against the RSC011 genome assembly by the software GMAP (version: 2024-02-22 with -max-intron-length-middle=10000 option) and then assembled intro transcripts by stringtie (version: v2.2.1) [22,23]. To annotate ORFs we modified the ppcac_generate_annotations.pl script of the PPCAC software to only consider ORFs with a starting Methionine. Raw RNA-seq reads from multiple developmental stages of *P. pacificus* strain RSC011 were assembled using the Trinity software (version: 2.2.0 with –normalize_reads option) [24]. Subsequently, the assembled transcripts were aligned against the polished RSC011 genome by GMAP. The resulting alignment file was taken as input to stringtie and was processed in the same way as the Iso-seq data. For visualization purposes, raw RNA-seq reads were also aligned against the polished RSC011 assembly by the software STAR (version: 2.7.10b with) and .bam files were created by samtools (version 0.1.18) [44,45].

### Genome assembly polishing

In order to reduce the number of problematic gene models, we employed whole genome resequencing data from more than 160 mutant lines in the RSC011 to screen for substitutions and indels that were called in more than 100 mutant lines [14]. This identified 1883 substitutions and 42,138 indels that were corrected in a new polished version of the RSC011 genome assembly (version 2) using the software mutsim [26].

### Gene annotation

During the course of this study, we generated three alternative versions of gene annotations for the new RSC011 genome assembly. In order to assess the effect of genome assembly polishing on gene annotation, we reran the original PPCAC gene annotation pipeline on the polished assembly using the same input data sets and parameters as described previously [14,15]. Specifically, these annotations were generated from transcriptomic evidence (ORFs in Trinity assembled transcripts) and homology data from the *P. pacificus* PS312 El Paco gene annotations (version 3) [17]. One representative gene model per 100-bp window was chosen by selecting the gene model with longest ORF while all other intersection gene models were then discarded [15]. This version is labeled as PPCAC1 polished (Table 1).

Next, we integrated the newly generated transcriptome data by providing them as additional input data into the PPCAC pipeline and prioritizing these gene models by adding a value of 100 to their ORF length. This version was labeled as PPCAC2 vanilla (Table 1) and formed the initial version for community curation. The final community curated version was labeled as PPCAC2 cream (Table 1).

### Community curation

For initial community curation, we defined problematic gene models as genes with more than three exons that were either annotated as 3’UTR or 5’UTR. After an initial assessment of hundreds of candidates by an experienced curator, three other community curators were then recruited from within the laboratory. These curators were then trained in a dedicated seminar where typical errors, evidence types, and solutions were presented and discussed.

The list of problematic gene models genes was then shared as an online spreadsheet across all community curators who could manually inspect the problematic locus in the genome browser (jbrowse on pristionchus.org) [25]. Different tracks in the genome browser included the alignment files for Iso-seq and RNA-seq data, as well as alternative gene models that were either created by stringtie or by exonerate. In addition, we labeled gene models based on the best BLASTP hit (e-value < 0.001) in *C. elegans* to visualize ORFs that are highly conserved. Using these data, the curators eventually proposed alternative gene models for a given candidate genes. Note that the curators were explicitly asked to skip problematic cases where different evidence types (RNA-seq alignments, gene orientation, homology information) did not point towards a clear solution. After the curation of the gene models with long UTRs, we implemented all suggested changes in an intermediate annotation and then scanned the whole genome at 20 kb resolution for any obvious annotation problems and proposed additional corrections. This also served to detect and correct any inconsistencies in the first round of community curation (e.g. addition of overlapping redundant gene models). After both rounds of curation, we compiled the final set of gene annotation and assigned new gene identifiers to the new gene models (S2 Table).

To facilitate comparisons between the previously published gene annotation (PPCAC1, Table 1) [14] and the current one (PPCAC2 cream, Table 1), we generated a mapping between gene models using an approach to identify syntenic orthologs between genomes [28].

## Supporting information

### S1 Table. Community curations

The table summarizes the updates to the PPCAC2 vanilla gene set by the community curators together with their classification.

### S2 Table. Gene Identifier conversion table

The table shows the mapping between the new gene identifier in the PPCAC2 cream gene set and the original sequence identifier.

### S3 Table. Gene Identifier mapping across versions

The table shows a mapping between gene identifiers of the PPCAC2 cream annotation and the previous annotation [14]. This mapping was generated by a method to identify syntenic orthologs between genomes [28].

## Data access

All transcriptomic data sets that were created for this study, have been submitted to the European Nucleotide Archive under the study accession PRJEB101809. The *P. pacificus* RSC011 genome assembly and gene annotations are also available on pristionchus.org.

## Acknowledgements

The authors thank the whole Sommer laboratory for helpful discussions. The authors would also like to thank Alejandro Sanchez-Flores and one anonymous reviewer for helpful comments on the manuscript.

## Author contributions

**Conceptualization:** Christian Rödelsperger.

**Data Curation:** Christian Rödelsperger.

**Formal Analysis:** Christian Rödelsperger.

**Funding Acquisition:** Ralf J. Sommer.

**Methodology:** Christian Rödelsperger.

**Investigation:** Christian Rödelsperger, Neha Agyal, Shiela Pearl Quiobe, Hao Wu, Waltraud Röseler, Dafne Ibarra-Morales.

**Project Administration:** Christian Rödelsperger, Ralf J. Sommer.

**Resources:** Ralf J. Sommer, Christian Rödelsperger.

**Software:** Christian Rödelsperger.

**Supervision:** Christian Rödelsperger, Ralf J. Sommer.

**Validation:** Christian Rödelsperger.

**Visualization:** Christian Rödelsperger.

**Writing – Original Draft Preparation:** Christian Rödelsperger.

**Writing – Review & Editing:** Ralf J. Sommer, Christian Rödelsperger.

